# Nasopharyngeal microbial communities of patients infected with SARS-COV-2 that developed COVID-19

**DOI:** 10.1101/2020.12.01.407486

**Authors:** Maria Paz Ventero, Rafael Ricardo Castro Cuadrat, Inmaculada Vidal, Bruno Gabriel N. Andrade, Carmen Molina-Pardines, Jose M. Haro-Moreno, Felipe H. Coutinho, Esperanza Merino, Luciana CA Regitano, Cynthia B Silveira, Haithem Afli, Mario López-Pérez, Juan Carlos Rodríguez

**Affiliations:** Microbiology Department, Alicante University General Hospital - Alicante Institute of Sanitary and Biomedical Research (ISABIAL), Alicante, Spain; Department of Molecular Epidemiology, German Institute of Human Nutrition Potsdam-Rehbrücke, Nuthetal, Germany; Embrapa Pecuária Sudeste, São Carlos, Brazil; Department of computer science, Cork Institute of Technology, Ireland; Evolutionary Genomics Group, División de Microbiología, Universidad Miguel Hernández, San Juan de Alicante, Spain; Infectious Diseases Unit, Alicante University General Hospital, Alicante Institute for Health and Biomedical Research (ISABIAL), Alicante, Spain; Department of Biology, University of Miami, Miami, USA

**Keywords:** COVID19, SARS-CoV-2, Microbiome

## Abstract

**Background:** SARS-CoV-2 is an RNA virus causing COVID-19. The clinical characteristics and epidemiology of COVID-19 have been extensively investigated, however studies focused on the patient’s microbiota are still lacking. In this study, we investigated the nasopharyngeal microbiome composition of patients who developed different severity levels of COVID-19. We performed Rdna-SSU (16S) sequencing from nasopharyngeal swab samples obtained from SARS-CoV-2 positive (56) and negative (18) patients in the province of Alicante (Spain) in their first visit to the hospital. Positive SARS-CoV-2 patients were observed and later categorized in mild (symptomatic without hospitalization), moderate (hospitalization) and severe (admission to ICU). We compared the microbiome diversity and OTU composition among severity groups using Similarity Percentage (SIMPER) analysis and Maaslin2. We also built bacterial co-abundance networks for each group using Fastpar.

**Results:** Statistical analysis indicated differences in the nasopharyngeal microbiome of COVID19 patients. 62 OTUs were found exclusively in SARS-CoV-2 positive patients, mostly classified as members of the phylum Bacteroidetes (18) and Firmicutes (25). OTUs classified as *Prevotella* were found to be significantly more abundant in patients that developed more severe COVID-19. Furthemore, co-abundance analysis indicated a loss of network complexity among samples from patients that later developed more severe symptoms.

**Conclusions:** Our preliminary study shows that the nasopharyngeal microbiome of COVID-19 patients showed differences in the composition of specific OTUs and complexity of co-abundance networks. These microbes with differential abundances among groups could serve as biomarkers for COVID-19 severity. Nevertheless, further studies with larger sample sizes should be conducted to validate these results.

**IMPORTANCE:** This work has studied the microbiota of the nasopharyngeal tract in COVID19 patients using advanced techniques of molecular microbiology. Diverse microorganisms, most of which are harmless or even beneficial to the host, colonize the nasopharyngeal tract. These microorganisms are the microbiota, and they are present in every people. However, changes in this microbiota could be related to different diseases as cancer, gastrointestinal pathologies or even COVID19. This study has been performed to investigate the microbiota from patients with COVID19, in order to determinate its implication in the pathology severity. The results obtained showed that it is possible that several specific microorganisms are present only in patients with severe COVID19. These data, could be used as a prognostic biomarker to early detect whose patients will develop a severe COVID19 and improve their clinical management.

## BACKGROUND

Severe Acute Respiratory Syndrome Coronavirus-2 (SARS-CoV-2) is a positive-sense single-stranded RNA virus causing Coronavirus Disease 2019 (COVID-19) [1]. On January 30, 2020, the World Health Organization (WHO) declared the COVID-19 outbreak as “public health emergency of international concern” and two months later on March 11^th^ as a pandemic. The SARS-CoV-2 virus was first reported in central city of Wuhan, Hubei province, China, and presented 70% of similarity with the SARS-CoV-1 virus [2] and 96% similarity with a bat coronavirus, which is an evidence of the original host of this zoonosis [1], although the exact source has yet to be elucidated. While the most common symptoms are fever, cough and dyspnoea, the disease can cause other less frequent clinical manifestations such as myalgia, headaches, breathlessness, fatigue and nausea [3].

Viruses and bacteria are often present in the respiratory tract of healthy and asymptomatic individuals [4]. Microaspiration of aerosols and direct mucosal dispersal is responsible for a constant inflow of microbes and viruses towards lower airways [4]. Disease and inflammatory processes that lead to the emergence of anaerobic zones, or mucus accumulation in the alveoli can drastically change the microbial community of the airways [4]. For example, in diseased individuals, the lung microbiome composition undergoes a decrease in diversity [7] accompanied by a shift in the dominant taxa: from Bacteroidetes to Gammaproteobacteria, a class that includes many respiratory pathogens.

Although the clinical characteristics and epidemiology of COVID-19 have been described [8,9,10], studies focused on the associations between the patient’s microbiota and the onset of the disease are still limited. This pilot study aims to characterize the nasopharyngeal mucosal microbial communities of SARS-CoV-2 infected patients. We investigated samples from a control group of SARS-CoV-2 negative patients and three groups of SARS-CoV-2 positive patients, divided according to disease severity: one group of symptomatic patients that did not require hospitalization, a second group of patients that were admitted to conventional hospitalization facilities, and a third group of patients that required admission to the ICU.

## METHODS

### Patients and experimental design

56 nasopharyngeal microbiome samples from SARS-CoV-2 positive patients and 18 samples from SARS-CoV-2 negative patients were collected during March and April of 2020 in the Emergency Service of Hospital General Universitario de Alicante (HGUA). Cobas SARS-CoV-2 PCR Test for the Cobas 6800 System (Roche Molecular Systems, Branchburg, NJ, USA) was used to detect the presence of SARS-CoV-2 [11].

Patients were first classified based on SARS-CoV-2 presence, and then regarding their later developments (hospital admission and severity). All samples were obtained before the onset of severe symptoms, and before any treatment was administered to the patients. Following these criteria, four groups were established: group 0: SARS-Cov-2 negative patients (n=18); group 1: mild COVID19 symptoms but no later hospital admission (n= 19); group 2: severe COVID19 symptoms followed by hospital admission (n=18); and group 3: patients with severe COVID19 symptoms which were eventually admitted into intensive care units (ICU) (n=19). Protocols were developed in accordance with the national ethical and legal standards, and following the guidelines established in the Declaration of Helsinki (2000). The research project was conducted under the written approval of the Ethic Committee of Clinical Research with Drug (In Spanish, CEIm) of the “Hospital General Universitario de Alicante (Spain)”, and in collaboration with the Biobank of Clinical and Biomedical Research Institute of Alicante (ISABIAL), which are included in the Valencian Network of Biobanks.

### DNA isolation and Sequencing

DNA from nasopharyngeal samples was isolated using the QIAamp DNA Mini Kit (QIAgen) following the protocol recommended by the manufacturer. Sequencing libraries were prepared according to the 16S Metagenomic Sequencing Library Preparation protocol distributed by Illumina. Briefly, the sequence spanning the hypervariable regions V3 and V4 of the 16S rRNA gene was amplified through PCR and amplicons were quantified using a Qubit 4 Fluorometer (Qubit dsDNA HS Assay Kit) and validated by 4200 TapeStation (company). Amplicons were sequenced with Illumina MiSeq System using the 2×300bp cartridge. The quality of raw sequences was assessed by FastQC software.

### Taxonomic classification of amplicon sequences

Paired end reads of 300 bp were generated with an average overlap of 140 bp. Sequences were trimmed using trimmomatic [12] and the resulting paired reads were merged using casper [13], generating individual fragments of about 460 bp. Given the uneven coverage between samples, the number of individual reads was standardized to 20,000 per sample, removing samples that did not reach this sequencing depth. Merged amplicon sequences were grouped in operational taxonomic units (OTUs) using cd-hit [14] with an identity of 97%. Sequences were queried against small subunits (16S) rRNA genes by the SILVA database [15] for taxonomic classification. Sequences with low identity (< 70%) to any reference 16S rRNA gene or classified as eukaryotic were excluded from further analysis.

### Testing for differences in taxonomic composition among patient groups

We sought to determine how different samples were grouped according to their OTU composition. To that end, non-metric multidimensional scaling (NMDS) analysis was performed based on Bray-Curtis dissimilarity measures were calculated among samples based on relative OTU abundances (i.e. percentages) through the Vegan (v 2.5-6) package in R (v 3.6.3). The relative abundances of OTUs were also used to test for statistically significant differences among severity groups. Group OTU compositions were compared through ANOSIM. Next, Similarity Percentage (SIMPER) analysis was used to determine which OTUs were responsible for driving the differences in community composition among groups. For this analysis, all six possible pairwise combinations of severity groups were tested.

### OTU association with COVID-19 severity

To infer associations between the severity of COVID-19 and the airways microbiome, general linear models (GLM) were built using the R package MaAsLin2 with centred log-transformed (CLR) OTUs counts as the dependent variable and the severity group (with group 0 and group 1 as references), adjusted by gender and age, as the independent variable. Only OTUs that presented a prevalence of 20% over the sample space were considered. The resulting p-values were adjusted for multiple testing using the Benjamini-Hochberg method (BH).

### Co-abundance networks for COVID-19 severity groups

Fastpar [16], a multi-thread implementation of the SparCC algorithm [17], was used to generate co-abundance networks among OTUs of each of the four severity groups with default parameters (50 iterations and correlation threshold of 0.2) and 1,000 bootstrap iterations to infer significance. Results were processed using an in-house ipython notebook to generate network matrices for visualization with Cytoscape 3.8 [18]. The network matrices were loaded in the Cytoscape 3.8 software, and connections filtered by p-value (≤ 0.05) and correlation (≤ −0.6 or ≥ 0.6).

## RESULTS

### Study Set

Seventy-four patients were included in this pilot study to assess associations between the nasopharyngeal microbiome composition and the severity of the COVID19 disease. However, only 65 samples remained after quality coverage control (see Material and Methods). Data including age, sex, diagnosis, hospital admission, and disease severity were registered (Table S1). Sixteen patients belonged to the negative control (Group 0, no-SARS-CoV-2), whereas the remaining patients were classified into three groups (Group 1, 2 and 3) according to the severity (see methods). The average age of the patients was *ca*. 60 years old and around 49% of them were diagnosed with pneumonia.

### Microbiome taxonomic composition differs among severity groups

The bacterial phylum Firmicutes was the most abundant in the nasopharynx microbiome among patients from all severity groups (52.94% ± 4.04%), followed by Bacteroidota (22.06% ± 6.07%), Proteobacteria (12.75% ± 7.28%) and Actinobacteria (5.4% ± 0.6%). At the genus level, *Streptococcus* was the most abundant taxon (25.23% ± 2.03%), followed by *Prevotella* (16.20% ± 5.66%), *Veillonella* (14.45% ± 2.20%), *Haemophilus* (5.28% ± 4.76%) and *Moraxello* (3.24% ± 3.6%) (Figure S1 and Table S2). A total of 62 OTUs were found exclusively in SARS-CoV-2 positive patients (at a minimum of three samples). Most of these OTUs were classified as members of the phylum Bacteroidetes (18) and Firmicutes (25). Notably, the most common genera among the OTUs found exclusively on COVID-19 positive patients were *Prevotella* (13), followed by *Leptotrichia* (4) and *Streptococcus* (4). Samples were compared based on the relative abundances of OTUs. This analysis revealed that samples did not cluster according to the severity group neither by hierarchical clustering (Figure S2A and 2B) or NMDS (Figure S2C). Nevertheless, the differences in OTU composition among severity groups were significant according to ANOSIM (R = 0.046, *p* = 0.036).

SIMPER analysis revealed that 25 OTUs were responsible for approximately 70% (p-value 0.04) of the differences in community composition between severity groups 1 and 3 (Table S3). These OTUs were classified as members of the phyla Bacteroidota, Firmicutes, Fusobacteriota and Proteobacteria. Eleven OTUs had higher average abundance among samples from severity group 1, among which were included three OTUs classified as members of the genus *Veillonella*. On the other hand, 14 OTUs were more abundant among samples from severity group 3, among which were included four OTUs classified as *Prevotella*.

### Multiple OTUs display differential abundance according to COVID-19 severity

Using group 0 as a reference, we identified a total of 10 significant associations between bacterial OTUs and patient severity (p-value < 0.05, q-value < 0.25), corrected for age and sex. Among those, 9 were positively associated (8 in group 2 and 1 in group 3 when contrasted with group 0) and 1 negatively associated (in group 3 contrasted with group 0) (Table S4, Figure 1A). Of the OTUs positively associated with severity, 3 were classified as members of the genus *Prevotella* (OTUs 4, 14 and 16). Due to the heterogeneity of group 0, we decided to investigate also the differences within the SARS-CoV-2 positive patients, using group 1 as reference. The GLM model showed just 1 significant OTU (OTU 16), a *Prevotella* also found to be significantly associated with severity in the first model (Table S4, Figure 1B). We did not find any OTUs significantly different between groups 1 and 0. Figure 1A shows the coefficients for all the significant OTUs found by both GLMs and figure 1B shows OTU 16 CLR transformed counts for all severity groups.

**Figure 1.**
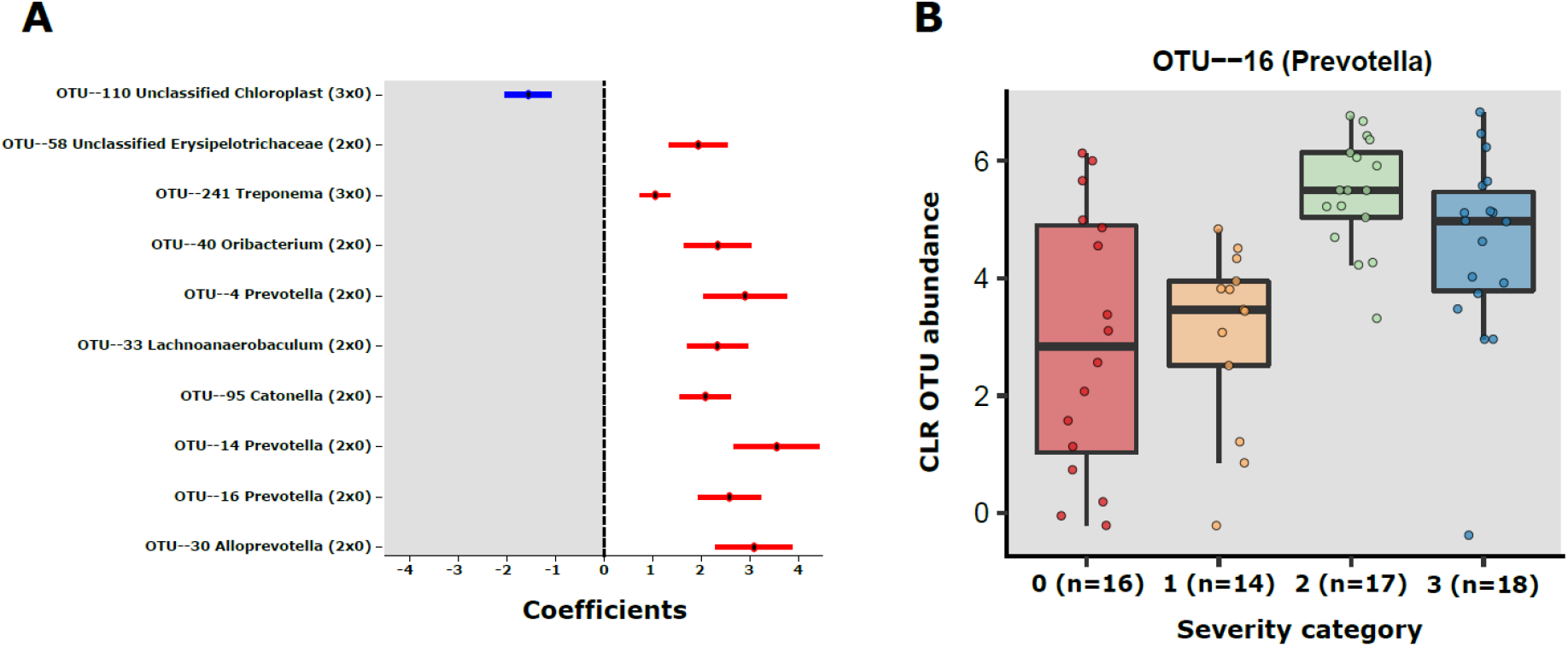
**A**) Error bar plot of the GLM coefficients interval for each OTU. The Y-axis shows the OTU number and Genus classification. The X-axis represents the CLR abundance. In red the positively associated OTUs and in blue the negatively associated OTUS. **B**) OTU 16 (*Prevotella*) center log transformed (CLR) abundance in the severity groups 0-3.

### Co-abundance networks for COVID-19 severity groups

In order to investigate how OTUs correlate in the different groups, we generated a total of 4 co-abundance networks, one for each severity group. For the severity group 0, the SARS-CoV-2 negative group, the network displayed 118 nodes with 179 edges. Regarding the other three severity groups, ranging from mild to high severity, the complexity of the network decreased with the increase of severity. The network for patients with mild symptoms (group 1) has 137 nodes with 457 edges, while the network for patients with severe symptoms but not admitted in ICU (group 2) had 129 nodes with 171 edges and the network for severe patients admitted in ICU (group 3) had 100 nodes and 148 edges. In the network of severity group 1, OTU 16 (*Prevotella*, associated with severity in two GLMs) displayed 18 co-abundant OTUs connected in the network in first degree (Figure 2). Among these connections, ten were negative associations while eight were positive. Most of these connections with OTU 16 were absent from networks of severity groups 2 and 3. Only 3 and 2 first degree connections remained in each of these networks respectively (Figure S3).

**Figure 2.**
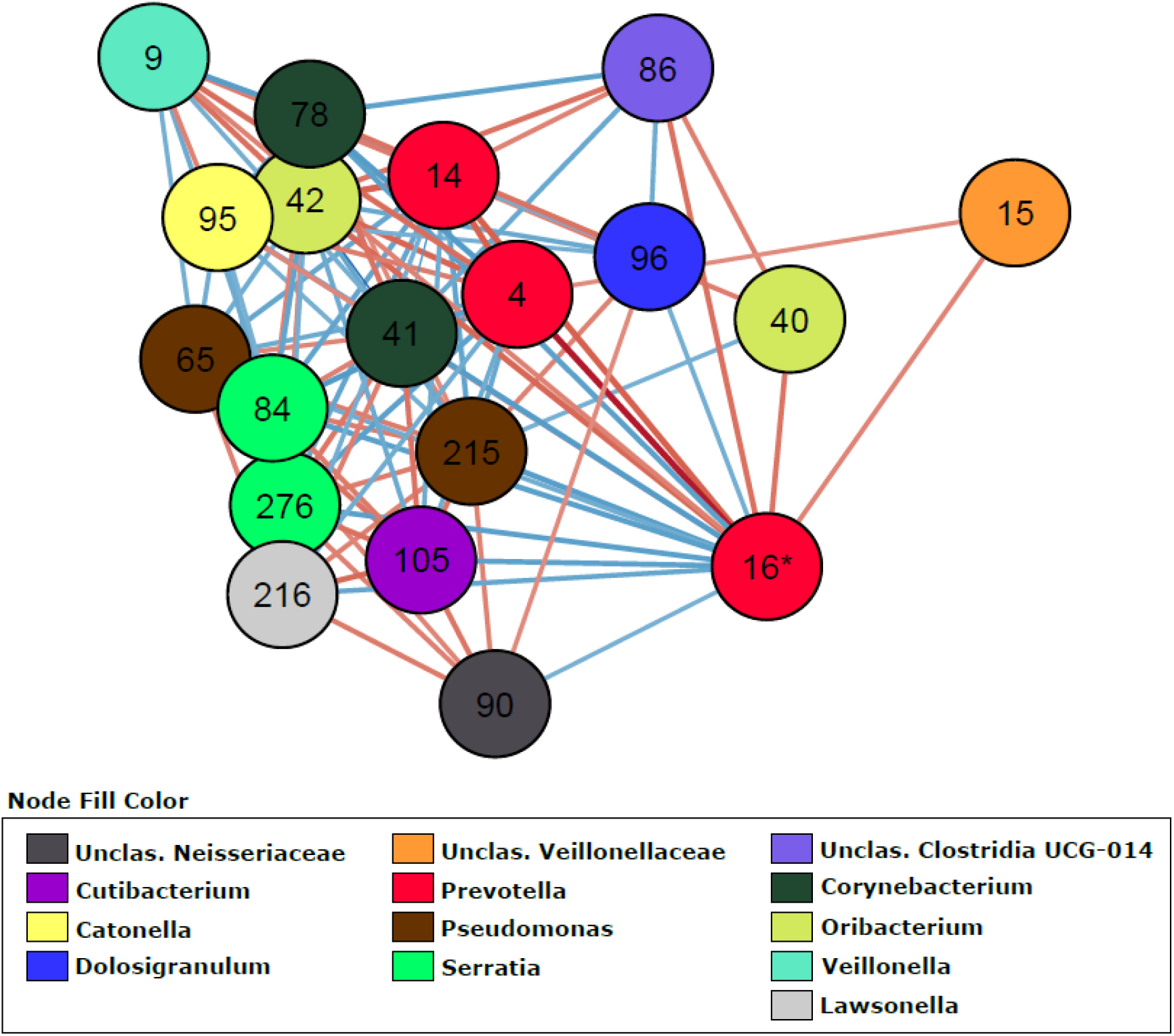
Co-abundance network (severity group 1) showing only first-degree neighbours of OTU 16 (*Prevotella* sp.). OTUs are represented by nodes and significant correlations by edges. Blue edges represent negative associations and red, positive associations. The colour of nodes was defined by the taxonomic classification of the OTU at Genus rank.

## DISCUSSION

In this preliminary study, analysis of the taxonomic composition of the samples showed differences between patients that developed different onsets of COVID-19. These changes in nasopharyngeal community composition are subtle, meaning that they are restricted to few taxa out of the complete meta-community. Nevertheless, there are detectable and significant changes among OTU abundances. These changes could be linked to the different severity groups, as we identified both taxa that were present exclusively among COVID-19 positive patients as well as those whose abundance was significantly higher or lower among different severity groups. Not only this, but also the complexity of co-abundance networks (which can be taken as a proxy for potential interactions between taxa), was decreased among patients that developed more severe cases of COVID-19. Below we discuss the mechanisms by which specific microbes might play a role in either enhancing or decreasing the severity of COVID-19. Those results suggest potential biomarkers for the onset of the disease.

### Potential associations between bacterial taxa and COVID-19 severity

Among the OTUs positively associated with COVID-19 severity, three were classified as members of the genus *Prevotella*, and one to a closely related genus, *Alloprevotella*. A recent study showed that *Prevotella* proteins can promote viral infection through multiple interactions with NF-κB signalling pathway, which is also involved in COVID-19 severity [19]. The genus *Prevotella* is usually considered commensal and, as such, rarely involved in infections. However, some strains have been identified as opportunistic pathogens in chronic infections, abscesses and anaerobic pneumonia [20,21,22,23]. The role of some strains of *Prevotella* in chronic mucosal inflammation has been demonstrated. They are involved with augmented T helper type 17 (Th17)-mediated mucosal inflammation, through activation of Toll-like receptor 2, followed by production of cytokines by antigen-presenting cells, including interleukin-23 (IL-23) and IL-1 [23]. The severe symptoms of COVID-19 are associated with cytokine storms, many of which are involved in TH17 type responses [24]. The significant association of *Prevotella* sp. and disease severity observed here suggests a possible link between *Prevotella* sp. and the COVID-19 through the activation of immunity signaling pathways that modulate inflammation, and this link should be further explored.

### Reduced network complexity among patients who later developed more severe COVID-19

Several studies demonstrated the usefulness of co-abundance networks to elucidate changes in the microbiome associated with human diseases [26,27,28,29]. By switching from individual OTU associations to a community interaction approach it is possible to attain a better understanding of the dynamic of microbiome/phenotype associations, revealing microbial consortia (and not only an OTU) that might be collectively influencing the host phenotype. Our linear models showed OTU 16 (*Prevotella* sp.) as an important OTU associated with severity. This OTU had the highest number of connections in the network, followed by OTU 9 (*Veillonella* sp.). Of the four networks generated, the severity group 1 network showed the higher number of interactions with this OTU. Ecological networking, *in vitro* and clinical studies showed that *Prevotella* sp. and *Veillonella* sp. are keystone species in microbiomes during airway disease progression, especially in diseases associated with mucus accumulation such as cystic fibrosis [30–32]. These anaerobes are efficient at degrading mucin molecules on the airway mucosa, releasing byproducts that enable the colonization and growth of pathogenic bacteria that are poor at degrading mucus for growth [33]. In COVID patients, *Prevotella* sp. and *Veillonella* sp. could have a similar role due to the decreased mucociliary clearance caused by the viral infection [34]. Lower rates of clearing increase the residence time of *Prevotella* sp. and *Veillonella* sp. in the airways, likely increasing their mucus metabolism and enabling further colonization by pathogenic bacteria that may cause pneumonia.

OTU 96, classified as *Dolosigranulum sp*., was identified in the group 1 network by having a negative relationship with OTU 16 (*Prevotella* sp.) as first-degree neighbor (Figure 3). OTU 96 did not pass the q-value threshold established for the GLMs but shows significant p-value (0.003 in the model comparing group 2 and group 0, and 0.02 in the model comparing group 2 and group 1 as reference). The only species currently described in this genus is *Dolosigranulum pigrum*, which is commonly found in the nasopharynx microbiome and is predicted to benefit the host through protection against pneumococcal colonization [35–36] and through protection against inflammation damage [37]. One study also found a lower abundance of *Dolosigranulum* in children with Influenza A Virus compared to healthy children [38]. In addition, a study reported that patients with their airway microbiota dominated by *Corynebacterium* and *Dolosigranulum* experienced the lowest rates of early loss of asthma control and have a longer time to develop at least 2 episodes [39]. We did not identify Corynebacterium directly connected to OTU16 (*Prevotella* sp.), but OTU 78, classified as Corynebacterium is positively associated with OTU 96 (0.7479, p-value 0.001) in the co-abundance network from group 1 (Figure 3), indicating that in asymptomatic patients those two taxa are forming a consortium that might protect from disease development. This “consortium” was also implicated in resistance to recurrent ear infections and it was proposed as a probiotic candidate for upper respiratory tract infections [40]. The reason that we did not have lower q-value in our GLM for those two taxa could be the lack of power due to the small size of our study. Thus, these associations warrant further investigation.

## LIMITATIONS

The major limitation of our study is the small sample size. With only about 15 samples per severity group it is difficult to find statistically significant associations between microbiome composition and disease severity. Nevertheless, this limitation is more likely to lead to false negatives than to false positives. We also cannot rule out confounding factors that might explain the differences between groups. Another important limitation is the fact that we performed amplicon rather than whole genome shotgun sequencing. This leads to three issues. First, some of the bacterial diversity is lost due to the fact that the selected primers do not amplify the entirety of bacterial diversity. Second, some genomes have more than a single copy of the 16S operon, which can lead to an overestimation of their abundance in the samples. Third, without metagenomes (and metagenome assembled genomes) we could not make inferences about the presence of virulence factors and other features of the genomes of the microbes in our samples. We resorted to 16S amplification because our non-invasive approach to collect samples yields low DNA amounts that are inadequate for sequencing. However, as far as we know, this is a unique pilot study in the field. The aim is to be able to transfer the first useful results to help clinical practice in the fight against the virus and to optimize all the protocols and analyses for a second analysis in which the sample size will be much larger. We are currently working on collecting more samples and optimizing protocols that will allow us to obtain whole genome shotgun sequencing from them.

## CONCLUSION

Our data provides preliminary evidence of significant differences in the composition of the upper airway microbiome according to COVID-19 severity, suggesting potential biomarkers of disease severity. While the richness indexes did not show significant differences among groups, specific taxa were significantly associated with disease development. We also demonstrated that the complexity of the co-abundance network is decreased in patients who came to develop severe cases of the disease, indicating that the interactions between the taxa are also relevant to this process. Further studies will be necessary to shed light on the molecular mechanisms that give rise to these associations. Finally, we make no claim that the differences in microbiome composition reported here are the cause of of COVID-19 severity. Nevertheless, the significant associations found between these variables suggests that the role of the microbiome on the onset of disease severity warrants further investigation.

## DECLARATIONS

### Ethics approval and consent to participate

This study has the written approval of the Ethic Committee of Clinical Research of Alicante University General Hospital (Ref. CEIm: PI2020-052). The samples used in this work are from clinical nasopharyngeal aspirates used to diagnose the COVID19 pathology during the first emergency state in Spain and stored in the Alicante University General Hospital Biobank. In this period, it was allowed to collected samples by Biobanks without obtaining the Informed Consent (Dictamen COVID19 D.20200327/2 CEI DGSP-CSISP).

### Consent for publication

Not applicable

### Availability of data and materials

Raw data was deposited to the National Center for Biotechnology Information Sequence Read Archive under BioProject accession number PRJNA673585.

All data generated during this study are included in this published article [and its supplementary information files].

### Authors’ contributions

JCR conceived the study. MPV, IV, CM and EM collected the data. RC, BA, CS, JHN and FH analysed the data. MPV, RC, MLP, FH, BA and JCR wrote the paper. All authors reviewed and approved the final version of the manuscript.

## Acknowledgments

Not applicable

## SUPPLEMENTARY MATERIAL

**Figure S1.**
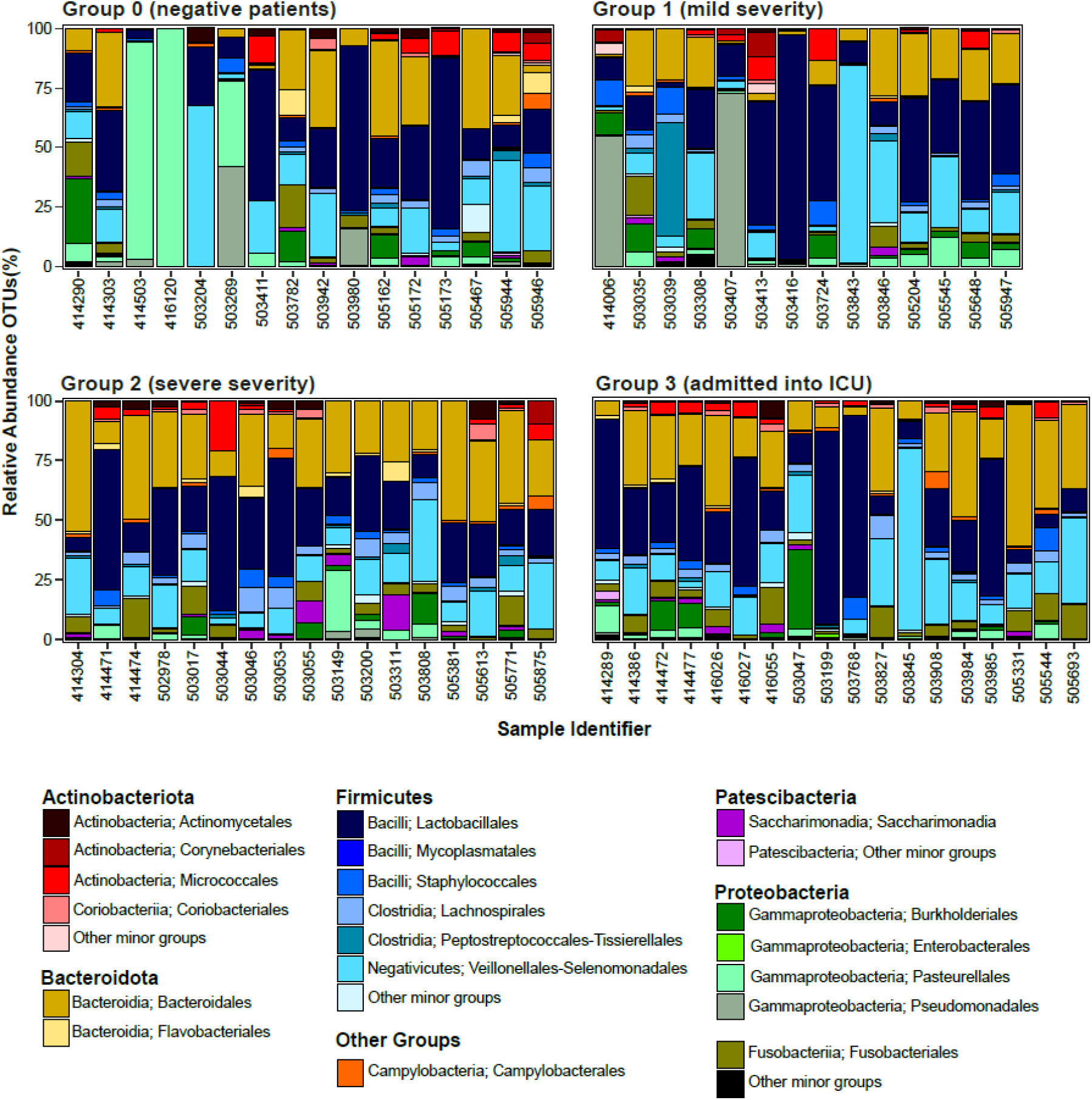
Relative abundance of bacterial populations, at genus level, in the microbiome of patients within COVID-19 severity groups. Only microorganisms with a relative abundance greater than 0.5% are shown in the legend.

**Figure S2.**
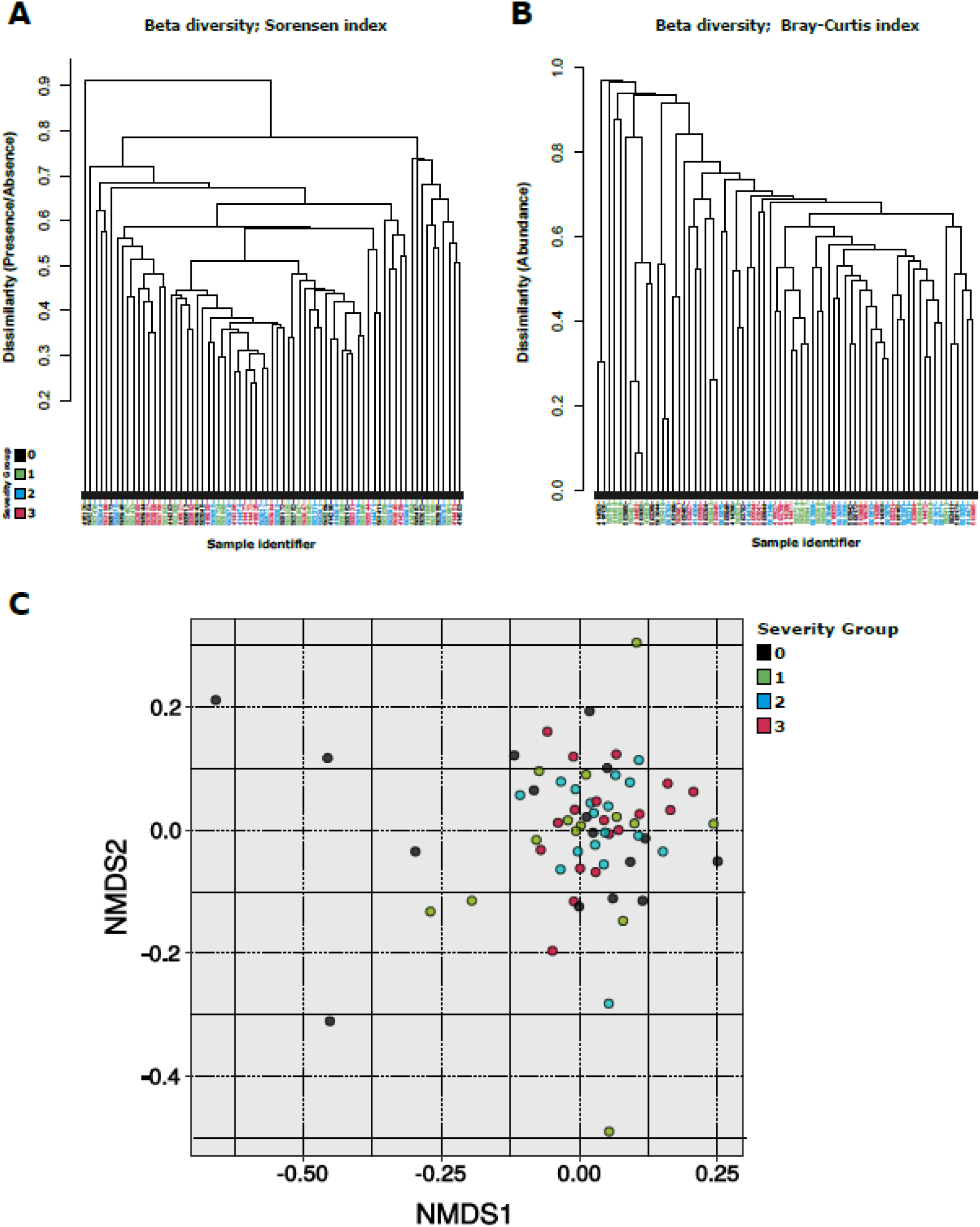
Beta diversity. Dendrogram based on **A**) Bray-Curtis dissimilarity and **B**) Sørensen dissimilarity values. **C) Comparison of sample taxonomic profiles by severity group.** Nonmetric multidimensional scaling was applied to determine the clustering patterns of samples according to their OTU abundance patterns. Each dot represents a sample color coded according to the severity group it belongs to. The closer the samples are, the more similar was their OTU abundance composition. No clear clustering of samples by severity group was observed.

**Figure S3.**
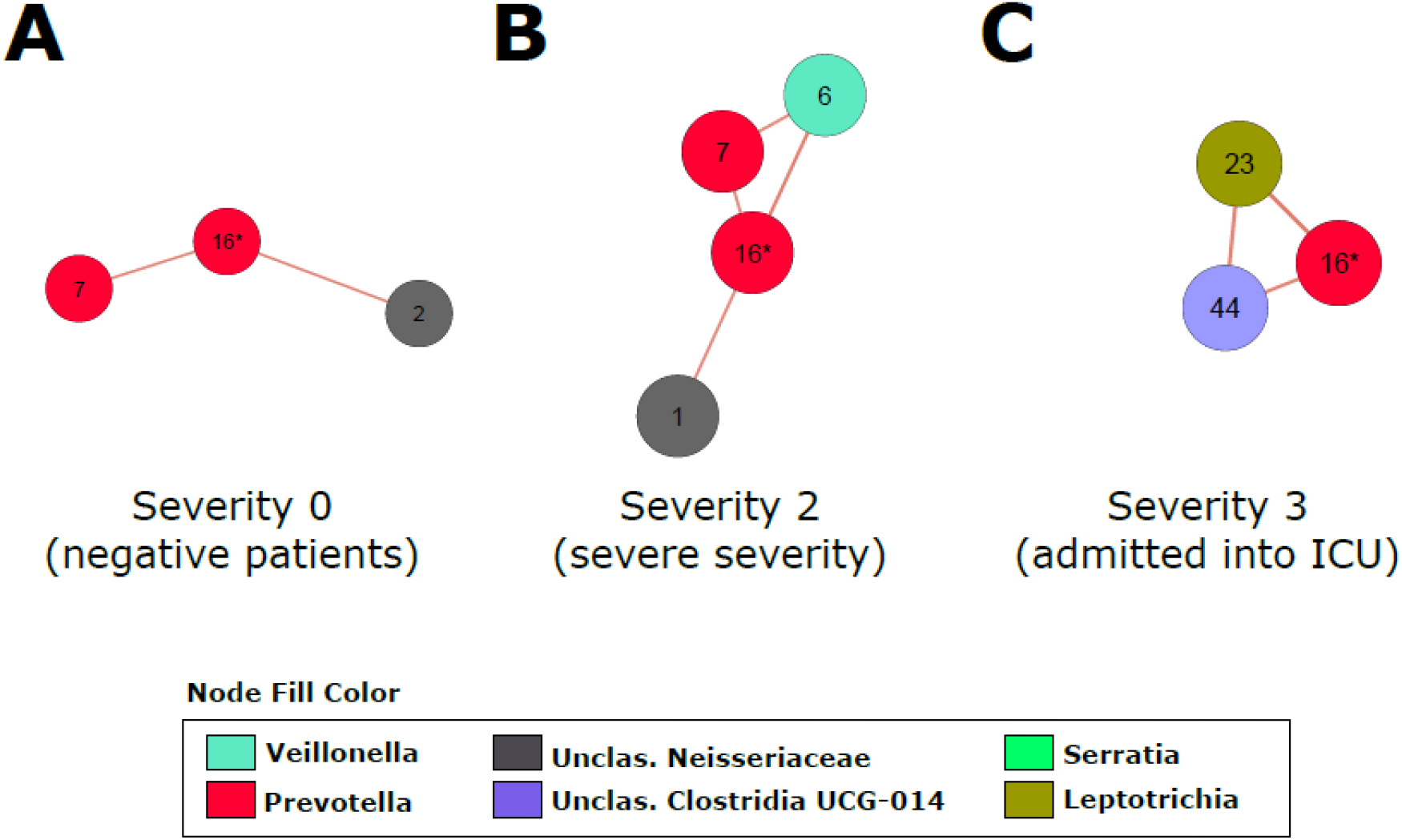
Co-abundance network showing only first-degree neighbours of OTU 16 (*Prevotella* sp.). **A)** Severity group 0 **B)** Severity group 2 and **C)** Severity group 3.

**Table S1**. Clinical features of patients.

**Table S2**. OTUs taxonomic classification.

**Table S3**. OTUs showing approximately 70% of the differences in community composition between severity groups 1 and 3 (SIMPER).

**Table S4.** Maaslin2 results (GLM) for OTUs associations (q-value < 0.25) for both models.

